# Signal complexity of human intracranial EEG tracks successful associative memory formation across individuals

**DOI:** 10.1101/180240

**Authors:** Timothy C. Sheehan, Vishnu Sreekumar, Sara K. Inati, Kareem A. Zaghloul

**Affiliations:** Surgical Neurology Branch, NINDS, National Institutes of Health, Bethesda, MD 20892; Office of the Clinical Director, NINDS, National Institutes of Health, Bethesda, MD 20892

## Abstract

Memory performance is highly variable between individuals. Most studies examining human memory, however, have largely focused on the neural correlates of successful memory formation within individuals, rather than the differences between them. As such, what gives rise to this variability is poorly understood. Here, we examined intracranial EEG (iEEG) recordings captured from 43 participants (23 male) implanted with subdural electrodes for seizure monitoring as they performed a paired-associates verbal memory task. We identified three separate but related signatures of neural activity that tracked differences in successful memory formation across individuals. High performing individuals consistently exhibited less broadband power, flatter power spectral density (PSD) slopes, and greater complexity in their iEEG signals. Furthermore, within individuals across three separate time scales ranging from seconds to days, successful recall was positively associated with these same metrics. Our data therefore suggest that memory ability across individuals can be indexed by increased neural signal complexity.

**Significance Statement:** We show that participants whose intracranial EEG exhibits less low frequency power, flatter power spectrums, and greater sample entropy overall are better able to memorize associations, and that the same metrics track fluctuations in memory performance across time within individuals. These metrics together signify greater neural signal complexity which may index the brain’s ability to flexibly engage with information and generate separable memory representations. Critically, the current set of results provide a unique window into the neural markers of individual differences in memory performance which have hitherto been underexplored.

## Introduction

Some people consistently have better memory than others. This variability in memory performance between individuals, and even within individuals from moment to moment, is quite familiar in our daily lives. In the study of memory, however, this variability has largely been viewed as a problem that needs to be addressed through proper experimental design. As a result, the neural mechanisms that give rise to such variability have been relatively unexplored. Understanding the source of such variability between individuals can provide valuable insights into how the brain is able to successfully form and retrieve memories.

Studies of memory have typically attempted to eliminate the variability in neural activity and memory performance between individuals by regressing it out. In many paradigms evaluating memory-related changes in oscillatory activity, for example, data within each individual are normalized so as to only examine relative changes in activity when events are either successfully remembered or forgotten, yielding what has been termed the subsequent memory effect (SME). Positive and negative SMEs have been reported in different frequency bands (Hanslmayr and Staudigl, 2014; Hanslmayr et al., 2012), yet how these effects should be properly interpreted has been problematic given conflicting reports of positive low frequency SMEs in some studies (Hanslmayr et al., 2011; Osipova et al., 2006; Sederberg et al., 2003) and negative low frequency SMEs in others (Fell et al., 2011; Guderian et al., 2009; Sederberg et al., 2006). Hence, normalized SMEs studied in isolated frequency bands may not provide a complete description of the neural correlates of memory. Moreover, these approaches have not addressed the larger question of how variability in neural activity may be related to variability in memory performance.

An alternative and complementary approach that has emerged in response to the conflicting SME data is to describe the changes in low and high frequency activity as arising from the same phenomenon, one that produces an overall change in the structure of the entire power spectral density (PSD) (Voytek et al., 2015). Spectral power decreases linearly with frequency on a log-log scale over a broad range of frequencies (Dehghani et al., 2010; He et al., 2010; He, 2014; Miller et al., 2009; Milstein et al., 2009). Importantly, neuronal activation results in a flatter PSD slope (He, 2011), reflecting decreases in lower frequency and increases in higher frequency power (Podvalny et al., 2015). These findings have led to the suggestion that flattening of the PSD slope, and the associated changes in spectral power, may therefore be a signature of increased asynchronous neuronal activity (Burke et al., 2015; Ray and Maunsell, 2011; Voytek and Knight, 2015).

Viewed from an information coding perspective, the PSD slope and oscillatory power of a neural signal, by indicating the extent of synchrony in the underlying neural activity, may be a proxy for neural signal complexity and underlying information content (Hanslmayr et al., 2012). Direct measures of complexity of neural signals such as sample entropy have supported this suggestion by demonstrating that more complex brain dynamics underlie enhanced cognitive performance (McIntosh et al., 2008), likely signifying a greater capacity to encode and process information. Indeed, several groups have advanced the notion that complexity in neural activity is functionally relevant and affords greater flexibility for cognitive processing (Deco et al., 2009, 2013; Faisal et al., 2008; Garrett et al., 2011, 2013; Grady and Garrett, 2014; MacDonald et al., 2006; Sleimen-Malkoun et al., 2015; Stein et al., 2005). Therefore, these metrics may together reflect a general capacity for processing information that may be particularly relevant for memory formation.

In this scenario, then, a possible explanation for the variability in memory performance between individuals is that different brains may exhibit differences in complexity, allowing a greater number of unique cognitive states that are relevant for encoding memories. We investigate this possibility here by examining changes in spectral power, PSD slope, and the sample entropy of neural signals captured from intracranial electrodes as participants perform a paired associates verbal episodic memory task. We were specifically interested in whether these metrics exhibit differences between individuals, and changes within individuals across time, that correlate with memory performance, and find that such measures of complexity and general information processing are indeed behaviorally relevant when forming and retrieving memories.

## Materials and Methods

### Participants

43 participants [23 male; age range 13-59; 32.1 ± 11.7 (mean ± SD) years old], with drug resistant epilepsy underwent a surgical procedure in which platinum recording contacts were implanted subdurally on the cortical surface as well as deep within the brain parenchyma. For each participant, the clinical team determined the placement of the contacts so as to best localize epileptogenic regions (see Figure 1C for electrode coverage). Preoperative clinical fMRI testing results were available for 37 participants, and 36 of these participants exhibited fMRI activity consistent with left language dominance. The Institutional Review Board (IRB) approved the research protocol, and informed consent was obtained from the participants or their guardians. Data from a subset of participants were initially collected and analyzed for previous publications (Yaffe et al., 2014, 2017; Greenberg et al., 2015; Haque et al., 2015). Computational analyses were performed using custom written MatLab (The MathWorks, Inc., Natick, MA) scripts.

**Figure 1.**
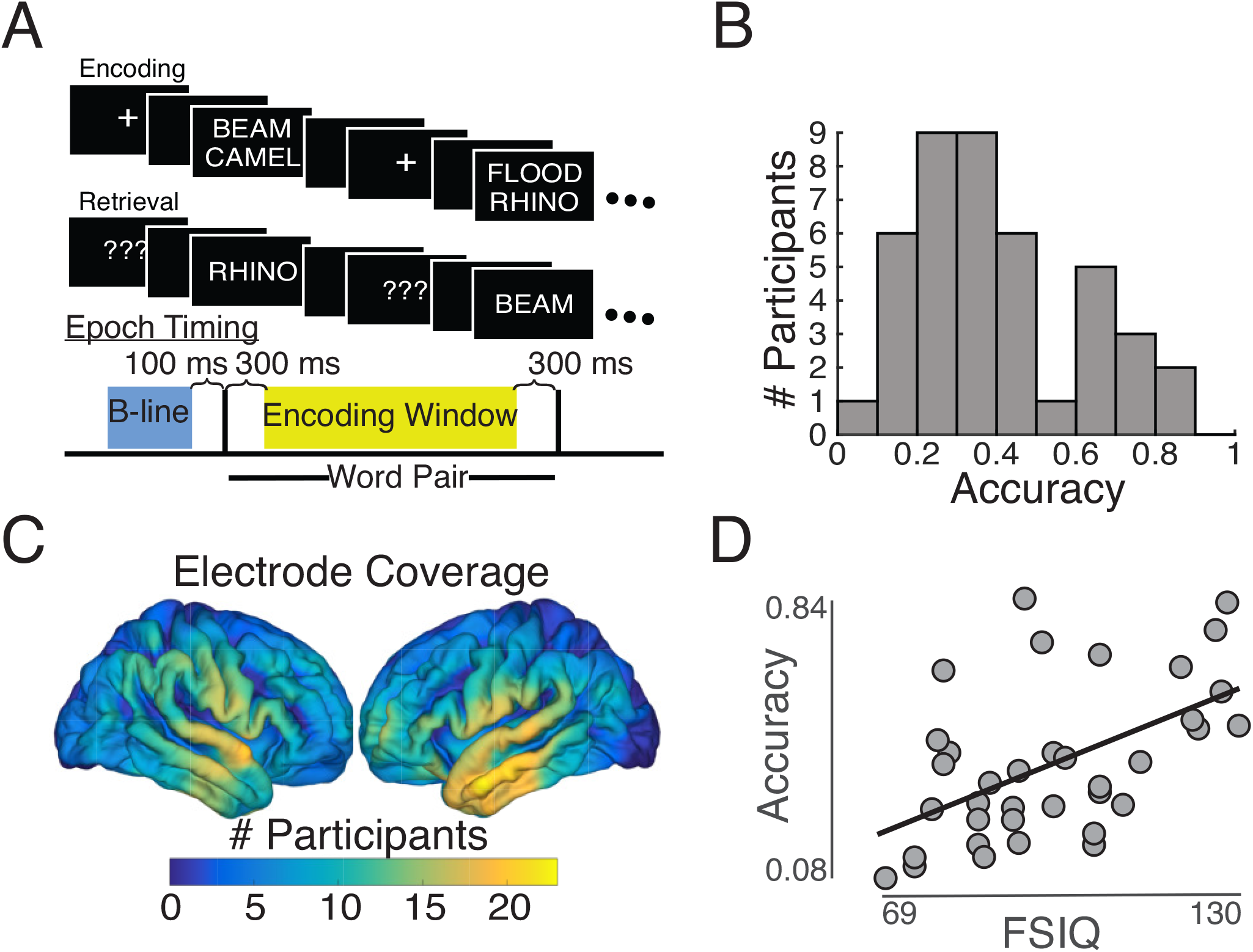
Paired associates task and subject distribution (*A*) Paired associates memory task schematic. (*B*) Average performance distribution across subjects, distribution is bimodal ranging between 0.05 and 0.84 with a median accuracy of 0.36 (N=43). (*C*) Electrode coverage by spatial region of interest. Colormap reflects number of electrodes within 12.5 mm. (*D*) Correlation between full-scale IQ and accuracy across subjects *r_s_* = 0.51, *p* = .0017, *N* = 35. Line is standard least squares regression line.

### Paired associates task

Each participant performed a paired associates verbal memory task (Figure 1A). In the task, participants were sequentially shown a list of word pairs (encoding period) and then later cued with one word from each pair selected at random (retrieval period), and were instructed to say the associated word into a microphone. Each participant performed one of two versions of the task that had slight differences in the experimental details. As the tasks did not differ in the fundamental objectives and performance was indistinguishable between groups, we combined the data from both sets of tasks for subsequent analyses.

A single experimental session for each participant consisted of 15 or 25 lists, where each list contained either four or six pairs of common nouns shown on the center of a laptop screen, depending on whether the participant completed the first or second version of the task respectively. Although different participants performed the task with different list lengths, the number of pairs in a list was kept constant for each participant. Words were chosen at random and without replacement from a pool of high-frequency nouns and were presented sequentially and appeared in capital letters at the center of the screen. Study word pairs were separated from their corresponding recall cue by a minimum lag of two study or test items. During the study period (encoding), each word pair was preceded by an orientation stimulus (either a ‘+’ or a row of capital X’s) that appeared on the screen for 250-300 ms followed by a blank interstimulus interval (ISI) between 500-750 ms. Word pairs were then presented stacked in the center of the screen for 2500 ms followed by a blank ISI of 1500 ms with a jitter of 75 ms in the first version of the task, or for 4000 ms followed by a blank ISI of 1000 ms in the second version. Following the presentation of the list of words pairs in the second version of the task, participants completed an arithmetic distractor task of the form A + B + C = ? for 20 seconds.

In both task versions, during the test period (retrieval), one word was randomly chosen from each of the presented pairs and presented in random order, and the participant was asked to recall the other word from the pair by vocalizing a response. Each cue word was preceded by an orientation stimulus (a row of question marks) that appeared on the screen for 250-300 ms followed by a blank ISI of 500-750 ms. Cue words were then presented on the screen for 3000 ms followed by a blank ISI of 4500 ms in the first version of the task, or for 4000 ms followed by a blank ISI of 1000 ms in the second version. Participants could vocalize their response any time during the recall period after cue presentation. We manually designated each recorded response as correct, intrusion, or pass. A response was designated as pass when no vocalization was made or when the participant vocalized the word ‘pass’. We defined all intrusion and pass trials as incorrect trials. A single experimental session contained 60, 100, or 150 total word pairs, or trials, depending on the task version. We included only participants who engaged in at least two separate sessions of the paired associates task such that each participant completed between 2-5 sessions taking 31.1 ± 1.7 (mean ± SEM) minutes each with a median of 24.8 hours in between sessions.

### Intracranial EEG (iEEG) recordings

Intracranial EEG (iEEG) signals were referenced to a common electrode and were resampled at 1000 Hz. We applied a fourth order 2 Hz stopband butterworth notch filter at 60 Hz to eliminate electrical line noise. The testing laptop sent synchronization pulses via an optical isolator into a pair of open lines on the clinical recording system to synchronize the iEEG recordings with behavioral events.

We collected electrophysiological data from a total of 3756 subdural and depth recording contacts (PMT Corporation, Chanhassen, MN; AdTech, Racine, WI). Subdural contacts were arranged in both grid and strip configurations with an inter-contact spacing of 5 or 10 mm. Contact localization was accomplished by co-registering the post-op CTs with the post-op MRIs using both FSL Brain Extraction Tool (BET) and FLIRT software packages and mapped to both MNI and Talairach space using an indirect stereotactic technique and OsiriX Imaging Software DICOM viewer package. The resulting contact locations were subsequently projected to the cortical surface of a population average brain. Pre-op MRI’s were used when post-op MRI images were were not available.

We took several steps to reduce the influence of pathologic activity on our results. First, we excluded from further analysis 414 electrodes identified clinically as having prominent interictal or ictal activity based on the evaluation of a board-certified epileptologist. To minimize the extent of transient epileptic activity (interictal discharges) in the remaining electrodes, we then performed an iterative cleaning procedure on the common averaged electrode signals to eliminate both electrodes and trials with a kurtosis greater than 2.8 or a variance greater than 2.2 standard deviations from the persistent sample’s mean. This procedure eliminated an additional 698 electrodes from further examination as well as 1047 out of 12650 individual trials. The remaining 2644 electrode contacts recorded over 11576 trials were used to generate our data set.

We analyzed iEEG data using bipolar referencing in order to reduce volume conduction and confounding interactions between adjacent electrodes (Nunez and Srinivasan, 2006). We defined the bipolar montage in our dataset based on the geometry of iEEG electrode arrangements. For every grid, strip, and depth probe, we isolated all pairs of contacts that were positioned immediately adjacent to one another; bipolar signals were then found by differencing the signals between each pair of immediately adjacent contacts. The resulting bipolar signals were treated as new virtual electrodes (referred to as electrodes throughout the text), originating from the mid-point between each contact pair. All subsequent analyses were performed using these derived bipolar signals. In total, our dataset consisted of 2347 bipolar referenced electrodes, derived from the set of original monopolar electrodes that remained following our cleaning procedure as described above.

### Data analyses and spectral power

We computed the spectral power for each bipolar electrode at every time point during the experimental session by convolving the raw iEEG signal with complex valued Morlet wavelets (wavelet number 6) to obtain the magnitude of the signal at each of 30 logarithmically spaced frequencies ranging from 3 to 180 Hz. We squared and log-transformed the magnitude of the continuous-time wavelet transform to generate a continuous measure of instantaneous power. During every trial, we convolved each wavelet with two separate time windows - a baseline period extending from 600 to 100 ms before word pair presentation, and an encoding period from 300 ms after word pair presentation until 300 ms before the offset of the word pair from the display screen (Figure 1A). In addition, we computed power for ten 2000 ms windows from the beginning of the clinical recording segment before task specific activity began for each session and averaged those to get an extra-task window. We included an additional 1000 ms buffer on either side of each time window to minimize any edge effects and which was not subsequently analyzed.

To examine the relation between overall raw power and performance across participants, we used the above measures of raw spectral power. To examine how changes in power on individual trials affected performance, we *z*-scored each sessions power values independently to remove the effects of across participant and session level variations.

### Calculating spectral slope

To understand how spectral power changes as a function of frequency, we calculated spectral slope. For each participant, we computed an average power spectral density (PSD) across all trials and electrodes and computed slope in log-log space across the broadband range of 10-100 Hz (Podvalny et al. (2015)). To identify the general 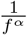 slope of the spectrum and avoid contamination of narrowband oscillations, we used a robust fitting algorithm with bisquare weighting (MATLAB *robustfit.m* function). Additionally, we computed slope over a range of frequency values and spectral widths as described in the Results. We defined spectral width using units of octaves such that the spectral width of a given slope was equal to the *log*_2_ of the ratio of the highest frequency to the lowest frequency.

### Calculating sample entropy

We used a metric of sample entropy to measure the complexity of the iEEG signal. Sample entropy, by construction, is a measure of predictability. Specifically the sample entropy (SampEn) of a time series is the negative natural logarithm of the conditional probability that any two sub-sequences of length m within the series, that are similar within a tolerance r, remain similar at length m+1 (Richman and Moorman, 2000). Two patterns that are close together in m-dimensional space and that remain close together in m+1-dimensional space indicate fewer irregularities or less complexity in the signal. Similarity is measured using the Chebyshev distance between the two sub-sequences. A smaller value of SampEn denotes greater repetitiveness and less complexity in a given signal.

For an embedding dimension m and a tolerance r, the formal equations for the calculation of sample entropy for a given time series of total length N are as follows (Sokunbi et al., 2013; Vakorin and McIntosh, 2012):

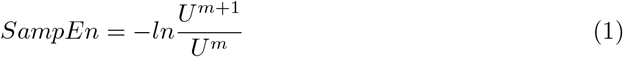

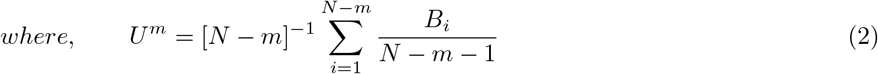

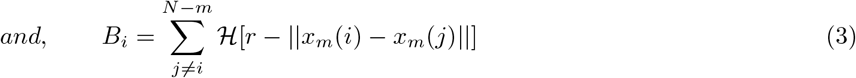

where ║ · ║ refers to the maximum norm, *x_m_*(*i*) is a vector {*x_i_*, *x*_*i*+1_,…,*x*_*i+m*−1_} within our time series, and 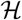 is the Heaviside step function. *B_i_* is the number of m-dimensional vectors that are within a tolerance of *r* from a given template *x_m_*(*i*), excluding self-matches. *B_i_* is normalized by the number of possible matches, *N – m –* 1, and averaged over the *N – m* possible template vectors to get *U^m^*, the probability of any two m dimensional vectors in a series being within a Chebyshev distance r.

An embedding dimension m of 2 and a tolerance *r* of 0.2\**std*(*x*(*t*)) were used in all analyses. Of note, the number of 3 element matching template sequences is necessarily less than or equal to the number of 2 element matching template sequences, implying that the ratio 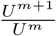 in Equation 1 is bounded between 0 and 1. Therefore, the range of SampEn is [0, ∞). For computational considerations, we down-sampled all iEEG signals to 250 Hz for this analysis, making our sampling period in between points in x 4 ms. We excluded the few trials with zero matching samples of length 3 to avoid infinite values.

### Commonality Analysis

In order to understand whether the metrics of power, spectral slope, and sample entropy uniquely account for variance in memory performance across participants or if they are redundant, we performed a commonality analysis (Nimon et al., 2008; Ray-Mukherjee et al., 2014) which partitions variance (*R*^2^) into parts that are unique to each predictor variable and those that are shared between all possible combinations of the predictors. n order to remain consistent with our rank based analyses used throughout the text and to remain sensitive to non-linear relationships, commonality analysis was performed on the ranks of our neural and performance measures. The unique contribution of a predictor is calculated as the proportion of variance attributed to it when it is entered last in a regression analysis. For example, consider a hypothetical case where dependent variable y is explained by two predictors *i* and *j*, the total variance in y explained jointly by both variables is 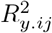, while the variance in *y* that is explained by *i* is 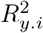, and the variance explained by *j* is 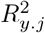. The unique contribution of a given variable is obtained by subtracting the contribution of the other variable from the joint contribution 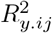. Therefore, the variance uniquely explained by *i* and *j* respectively are:

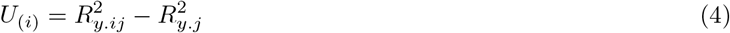

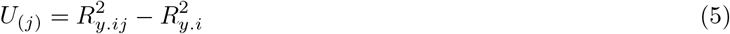

The common variance in y explained by *i* and *j* is equal to the total variance explained jointly less the unique contributions of *i* and *j*:

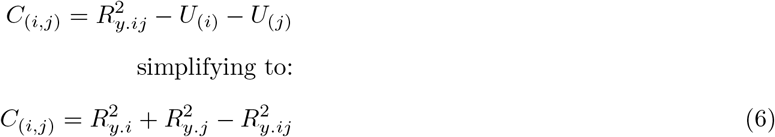

Commonality analysis decomposes explained variance into 2^*k*^ – 1 independent effects for *k* predictor variables. Therefore the number of effects increases exponentially with the number of predictors. We used the R package *yhat* (Nimon et al., 2013) to perform commonality analysis.

### Anatomic visualization

To visualize how the relation between spectral power and task performance is spatially distributed, we created 1441 regions of interest (ROI) evenly spaced across a 1 cm x 1 cm grid covering the pial surface of a population average brain. In each participant, we identified all electrodes located within 12.5 mm of each ROI. We designated the raw power for each ROI in each participant as the average raw power across all electrodes assigned to that ROI. For each ROI that included electrodes from at least six participants, we determined the Spearman’s correlation between raw power and task performance across the participants with electrodes contributing to that ROI. We therefore generated a value for the correlation between raw power and task performance for each ROI. Any ROI that contained electrodes from fewer than six participants was excluded from statistical analyses.

We generated cortical topographic plots of the anatomic distribution of these correlations by assigning each vertex in the 3D rendered image of the standard brain a weighted average of the mean value of each ROI that includes that vertex. Weighted values for each vertex were assigned by convolving a three dimensional Gaussian kernel (radius = 12.5 mm; *σ* =4.17 mm) with center weight 1 with the values of surrounding ROIs. We projected these vertex values onto the standard brain. Intensity varied as a function of the statistic metric in question, either Fisher-transformed correlation or *t*-score, in each ROI and with the standard deviation of the Gaussian kernel, which was used purely as a visualization technique.

### Statistical analysis

All statistical tests were assessed for significance using two-tailed distributions. As most of our distributions, including accuracy and raw power were not normally distributed, we utilized Spearman’s rank correlation when evaluating the monotonic relationship between two variables. Spearman’s correlation utilizes only the order of data points and is thus not biased by outliers as with Pearson’s correlation. We made an exception, however, when examining the relation between sessions within individual participants. Because we analyzed session counts as low as three, Spearman’s correlation is prone to produce extreme values of ±1 which cannot be analyzed with cohort level statistics, necessitating the use of Pearson’s correlation in this instance.

To compare correlations across participants, we used a Fisher *z*-transformation on the correlation coefficients calculated for each participant. The transformation stabilizes the variance of these correlations, reduces bias towards lower correlations, and results in a normalized distribution of coefficients. For each correlation, we therefore calculated the Fisher *z*-transform: 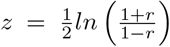 where *r* is the correlation coefficient. We utilized the mathematically equivalent formula, z = *arctanh*(*r*) in our calculations.

To determine whether any anatomic region exhibited a significant correlation across participants, we used a nonparametric spatial clustering procedure (Maris and Oostenveld, 2007). This procedure identifies contiguous ROIs where the distribution of correlation coefficients across participants significantly deviates from chance correlation while controlling for the family-wise error rate. Briefly, for each ROI, we calculated the true Fisher-transformed correlation coefficient between memory performance and raw spectral power across participants. We then generated 1000 permuted values for each ROI. In each permutation, we randomly assigned each participant a level of task performance drawn from the original distribution of task performance across participants without replacement. In this manner, each permutation involves a random pairing between task performance and raw spectral power. We then determined a *z*-score for each true value and each permuted value in each ROI by comparing that value to the distribution of permuted values. For the true data and for each permutation, we identified contiguous spatial clusters of ROIs, exhibiting *z*-scores with a magnitude greater than 1.96 (corresponding to a two-tailed *p*-value less than 0.05). For each cluster, we computed the cluster statistic as the sum of all *z*-scores in that cluster. In this manner, large magnitude cluster statistics can arise from large deviations in the distributions of correlation coefficients across participants extending over a small spatial region, or moderate deviations that extend over larger regions. We then calculated the exact two-tailed *p*-value for each cluster observed in the true dataset by comparing its cluster statistic to the distribution of largest cluster statistics drawn from each permutation. Clusters were determined to be significant and corrected for multiple comparisons if their *p* value calculated in this manner was less than 0.05.

To assess whether the relation between sample entropy and performance at different time scales was significantly from zero when summarizing across participants, we used a similar permutation procedure. In this case, for every ROI, we used a two-tailed *t*-test to compare the distribution of values to zero. This generates a *t*-statistic for the true data. Then, during every permutation, we randomly inverted the sign of the metric and produced a permuted distribution of *t*-statistics. We compared the true *t*-statistic to the permuted distribution to generate a *p*-value and *z*-score for every ROI. As above, we used a clustering procedure to identify contiguous ROIs with *p* < 0.05, assigned each contiguous cluster a cluster statistic based on the sum of the corresponding *t*-statistics, and then calculated the exact two-tailed *p*-value for each cluster observed in the true dataset by comparing its cluster statistic to the distribution of largest cluster statistics drawn from each permutation.

### Manual Inspection for Artifacts

To evaluate the influence of pathological activity on our results, a board-certified clinical epileptologist evaluated a subset of our recordings for the presence of interictal epileptiform discharges (IEDs), allowing us to examine the effects of this pathological activity on theta power in a given electrode or during a given trial, as well as on the average theta power for a given participant. For each participant, we selected and analyzed the five electrodes and ten trials exhibiting both the highest and the lowest magnitude theta power. Using custom viewing software, and blinded to the method of selecting trials or electrodes, the epileptologist was asked to evaluate whether a given trial did or did not contain epileptiform activity, and subsequently to identify the number of IEDs of any amplitude present in a specified bipolar electrode channel in a given two minute sample. To determine if IEDs were more likely during high theta power events or trials, we compared the two groups within each participant. There was no significant difference in the number of IEDs observed in a two minute period between low, 1.12 ± 1.39, and high 0.88 ± 1.72 theta power electrodes across ten participants (*t*(9) = 0.707, *p* = 0.50, *paired* t – test). There was also no significant difference between the percent of events exhibiting IEDs anywhere between low 45.0 ± 31.0 and high 61.0 ± 31.8 theta power trials *t*(9) = –0.97, *p* = 0.36. Lastly, to determine if IEDs were biased with respect to average power for each participant, we correlated total number of IEDs in our examined electrodes with average theta power and found there was not a significant correlation (*r_s_* = 0.40, *p* = 0.26, *N* = 10).

## Results

43 participants with drug resistant epilepsy who underwent surgery for placement of intracranial electrodes for seizure monitoring participated in a verbal paired associates task (Figure 1A). Participants studied 294.2 ± 20.0 (mean ± SEM) word pairs, split across multiple experimental sessions, and successfully recalled 40.1 ± 3.2% (mean ± SEM) words with a mean response time of 1837 ± 65 ms. Response accuracy across participants exhibited a bimodal distribution (Figure 1B). On 14.9 ± 1.7% of trials, participants responded with an incorrect word (intrusions) with a mean response time of 2687 ± 83 ms. For the remaining 44.9 ± 2.6% of trials, participants either made no response to the cue word, or vocalized the word ‘pass’ with a mean response time of 3494 ± 176 ms. We designated all trials in which a participant successfully vocalized the correct word as correct, and all other trials as incorrect. Recordings were included from all electrode contacts (number of participants with contacts in each cortical location shown in Figure 1C).

We measured full scale IQ (FSIQ) in 35 participants before electrode implantation as part of the routine clinical pre-operative evaluation. Participants had an average pre-operative FSIQ of 98.5 ± 2.9 (mean ± SEM). Across all sessions for each participant, we found that preoperative FSIQ significantly correlated with accuracy during the task *r_s_* = 0.51, *p* = .0017, *N* = 35; Figure 1D), suggesting that task performance is related to normal psychometric measurements.

### Raw power is negatively correlated with performance

Raw intracranial EEG (iEEG) power can reflect the extent of overall neural activity in each participant’s brain and has occasionally been shown to relate to a participant’s abilities (Hanslmayr et al., 2007). We were therefore interested in examining whether the raw overall power in each participant as captured by iEEG was related to their task performance. As typical spectral analysis involves examining changes in *z*-scored power relative to an individual’s baseline activity, this relation between raw power and task performance would be unexplored in most planned analyses.

In each participant, we extracted the raw spectral power contained in the signal during a baseline time window before word pair presentation and during the encoding period. To generate an overall level of broadband power for each participant, we averaged the extracted spectral power over all frequencies between 3 and 180 Hz (broadband power), over all trials, and over all electrodes for each time window in each participant. We found that the average raw broadband power during the encoding period demonstrated a significant negative correlation with accuracy during the task *r_s_* = –0.39, *p* = .01109, *N* = 43; Figure 2A). This was unchanged if we *z*-scored each frequency band across subjects to equalize contributions across bands. As with task performance, broadband power was also negatively correlated with with FSIQ across participants (*r_s_* = –0.408, *p* = .0348, *N* = 35).

**Figure 2.**
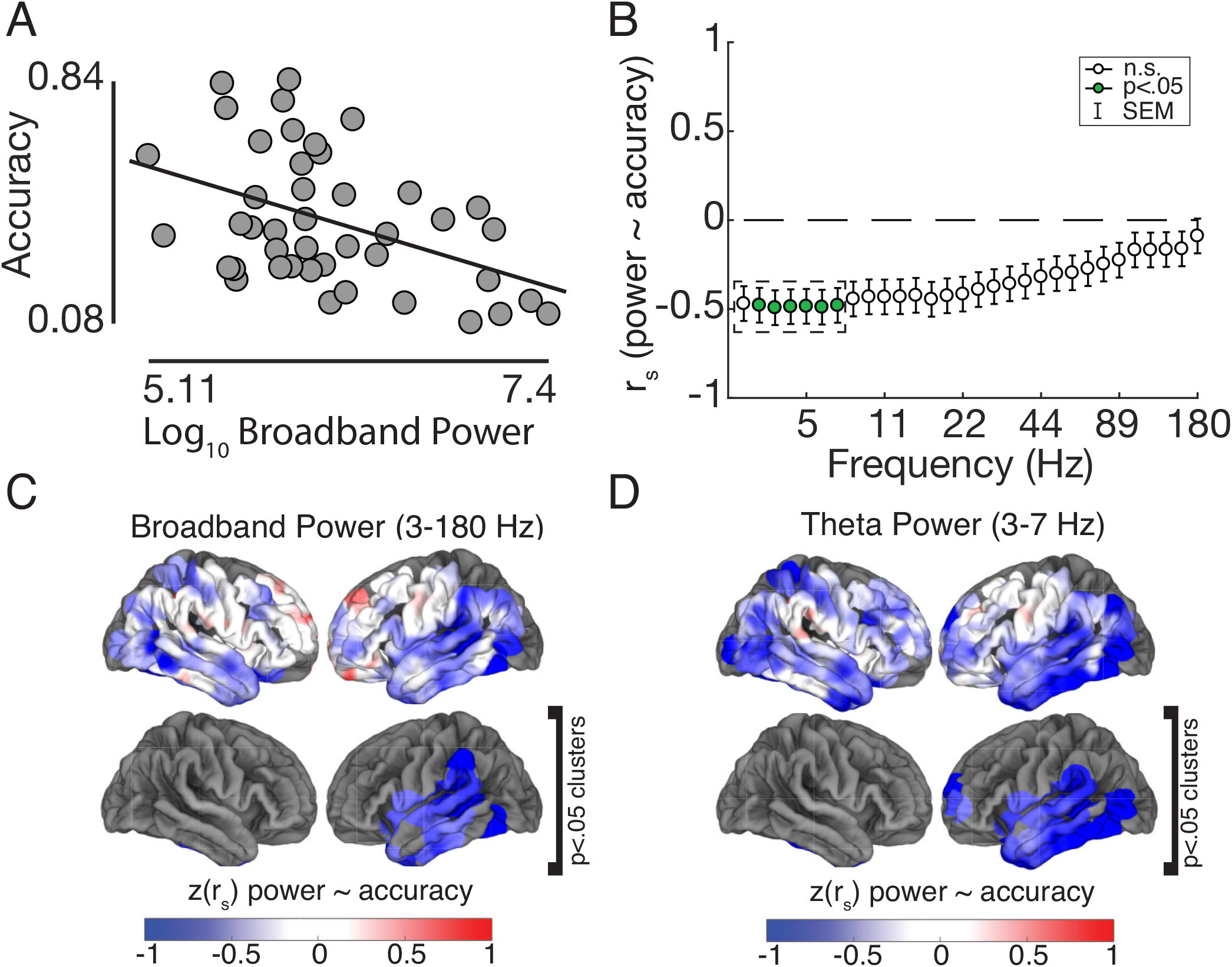
Baseline Power and Performance. (*A*) Average *log*_10_ broadband power across all trials and electrodes, range 5.11 to 7.40 (arbitrary units) is negatively correlated with performance *r_s_* = −.39, *p* = .011, *N* = 43. Line is standard least squares regression line. (*B*) Power ^~^accuracy correlation by frequency band. The negative correlation between power and accuracy exists across all bands and is significantly negatively correlated at all frequency bands between 3.5 to 9 Hz (*p* < .05, Bonferroni correct for 30 frequency bands). The error bars indicate standard error of the mean for Spearman’s correlation 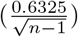. Theta power spectral region of interest is inside of dashed box. (*C*) Broadband (fisher transformed) correlation across spatial ROIs. Lower panel shows regions significant (*p* < .05) compared to a permuted distribution through a clustering procedure. (*D*) Same as *C* for theta band power.

We found that this relation between raw overall broadband power and task performance was robust and independent of the specific task periods. For example, raw broadband power during the baseline period before word presentation was also inversely correlated with task accuracy (*r_s_* = –0.38, *p* = .0125, *N* = 43). Moreover, we also found a significant relation between raw broadband power and task accuracy when we examined power separately during only correct or only incorrect trials (*r_s_* = –0.38, *p* = .0132 and *r_s_* = –0.38, *p* = .0114, respectively), suggesting that this effect reflects each participant’s overall baseline neural activity rather than simply the proportion of trials that featured successful encoding in each participant. Finally, to determine if this relation reflects each participants underlying physiology or is dependent on a task evoked state, we also examined this relation during an epoch recorded prior to the beginning of the task when the participant was awake, at rest, and under no instruction. We found the negative correlation between overall broadband power and task performance was also preserved during this extra-task period, suggesting that this effect is not task dependent but is related to baseline cognitive behavior (*r_s_* = –0.36, *p* = .0198). This finding departs from most memory studies in that we claim that our result does not depend on the fact that the subject is undertaking a memory task at the time, allowing us to generalize our electrophysiological correlates to normal daily activities.

We next examined whether the inverse correlation between raw power and task performance was specific to individual frequency bands by separately computing correlations between narrow band frequencies and task performance (Figure 2B). We found power at every frequency band between 3 and 180 Hz was negatively correlated with performance. All frequencies between 3.5 and 9 Hz had a significant negative correlation between overall raw power and accuracy when corrected for multiple comparisons across frequencies (Figure 2B, *p* < .05, Bonferroni corrected for 30 frequency bands). This suggests that this effect is spectrally broad but driven by low frequency activity. We therefore restricted subsequent power analyses to power averaged across the theta band which had a correlation of *r_s_* = –0.50, *p* = .0008 (3-7 Hz; Figure 2B, dashed box). As performance was shown to be strongly correlated with IQ, it is not immediately clear if the relationship between power and accuracy is simply a manifestation of a relationship between between power and general ability or if power explains additional variance in task performance not explained by IQ. While theta power is negatively correlated with IQ (*r_s_* = –0.44, *p* = .0074, *N* = 35), it is also correlated with performance after regressing out the variance in accuracy explained by IQ (*r_s_* = –0.42, *p* = .014, N=35), indicating it is independently predictive of task specific performance.

We were also interested in whether the relation between raw power and task accuracy varied across brain regions (regions of interest, ROIs; see Materials and Methods). For every ROI, we determined the correlation between both average raw broadband and theta power in all electrodes within that ROI and task performance across participants. The inverse correlation that we found between cortically distributed raw power and task performance localized to regions of the temporal and parietal lobes in both hemispheres (Figure 2C,D, *top*). Using a non-parametric clustering algorithm, we found that spatially contiguous regions exhibited a significant correlation across participants within the left temporal lobe for both broadband and theta band power (*p* < .05, permutation procedure; see Materials and Methods; Figure 2C,D, *bottom*). These data suggest that individuals with less broadband and low frequency power in the temporal lobe have greater ability to encode associative memories.

### Assessing cortical activation through PSD slope

The power spectral density (PSD) of iEEG signals falls off with frequency following a power law distribution. The slope of the PSD in log-log space has been shown to flatten in response to task activation (Podvalny et al., 2015), and the extent to which the slope flattens has been related to cognitive effort (Churchill et al., 2016). As such, we examined the overall raw PSD in each participant to determine whether the observed changes in broadband power with task performance may be related to changes in the slope of the PSD.

We first divided the participants into three terciles based on task performance (low, medium, or high accuracy) to visualize the average raw PSD in each cohort (Figure 3A). Below 30 Hz, the average PSDs of the three populations easily separate, with the lowest performing participants exhibiting the largest low frequency raw power. As suggested by our analysis examining the correlation between raw power and task performance, dividing participants into these terciles yielded a significant effect of performance tercile on low frequency power (ANOVA using average raw power < 30 Hz; *F*(2, 40) = 4.40, *p* = .019). At higher frequencies (> 30 Hz), however, the distinction between participant groups was negligible (*F*(2,40) = 0.913, *p* = .409).

**Figure 3.**
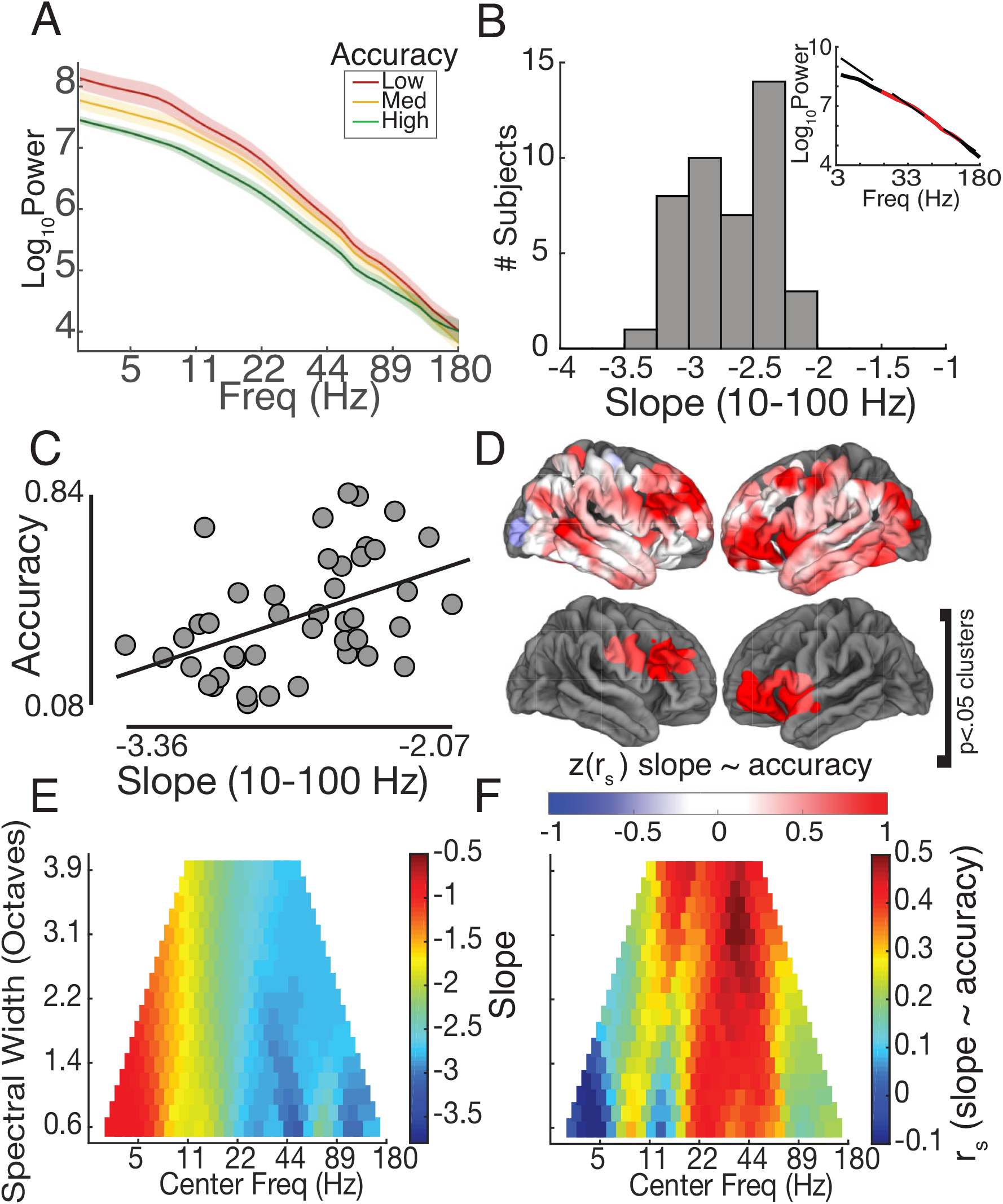
Spectral Slope and Performance. (*A*) Average power spectral density across tertials of subjects sorted by performance, shading shows standard error of the mean. (*B*) Distribution of average spectral slopes across subjects 2.67 ± .05 (mean ± SEM). Insert shows example subject, red is range of frequencies slope is calculated over (10-100 Hz) and dashed line shows robust fit line. (*C*) Spectral slope is positively correlated with accuracy across subjects *r_s_* = 0.48, *p* = .0014, *N* = 43. Line is standard least squares regression line. (*D*) Correlation of spectral slope and accuracy across spatial ROIs. Lower panel shows regions significant (*p* < .05) compared to a permuted distribution through a clustering procedure. (*E*) Average spectral slope as a function of center frequency and spectral width. (*F*) Average correlation as a function of center frequency and spectral width as in (*E*).

We next calculated the slope of the average PSD in log-log space between 10-100 Hz for each participant (Figure 3B, *insert*). We chose this frequency range to avoid the low frequency knee and the effects of action potential contamination at higher frequencies (Podvalny et al., 2015). Across participants, PSD slope [range −3.36 to −2.07; −2.67 ± .05 (mean ± SEM) Figure 3B)] are in the range of those reported by those using similar metrics (Podvalny et al., 2015). Across participants, we found that PSD slope was positively correlated with task performance, such that participants with flatter slopes performed better (*r_s_* = 0.48, *p* = .0014, *N* = 43; Figure 3C). Slope was not significantly correlated with IQ (*r_s_* = 0.17, *p* = .32), and was still correlated with performance after regressing out the effects of IQ (*r_s_* = 0.49, *p* = .0032, *N* = 35). We examined the anatomic regions that demonstrated a significant relation between PSD slope and task performance (*p* < .05, permutation procedure) and localized them to the left and right frontal lobes (Figure 3D).

As with raw broadband and theta power, this relation between PSD slope and task performance was robust and independent of when during the task the calculation of PSD was made. We found participants with greater task performance had flatter slopes when examining recordings from the baseline period (*r_s_* = 0.45, *p* = .0026), during correct trials only (*r_s_* = .47, *p* = .0018), or during incorrect trials only (*r_s_* = 0.47, *p* = .0015). We found that like broadband power, the significant relationship between slope and accuracy was preserved when examining extra-task epochs during which participants were awake and at rest (*r_s_* = 0.44, *p* = .0034), indicating that as with raw power, this relation is not task evoked.

Although several studies have identified measures of broadband power or spectral slope as a proxy for spike rate (Manning et al., 2009), cortical activation (Podvalny et al., 2015), or the balance between cortical excitation and inhibition (Gao et al., 2017), there still remains no consensus regarding the frequency range over which one should calculate the PSD slope in order to identify the non-oscillatory components of spectral power. In our initial analysis, we used a range of 10-100 Hz. However, other groups have used different frequency ranges and it is possible that our findings are sensitive to this parameter.

To ensure that the observed relation between PSD slope and task performance was not specific to the range of frequencies we used to calculate PSD slope, we iterated through a library of different frequency windows, each comprised of a center frequency and a spectral width, to compute PSD slopes. We examined PSD slope using every possible frequency window between 3 and 180 Hz. We found that the slope was largely unaffected by spectral width, and that although the slope increased as a function of center frequency, beyond a center frequency of 20 Hz, average slope stabilized to an overall mean across participants of approximately 2.7 (Figure 3E). Above 18 Hz, the PSD slopes ranged between −2 and −3, while at lower center frequencies we observed PSD slopes as flat as −1. We examined how varying our measure of PSD slope affected the relation between PSD slope and task performance. We correlated each PSD slope calculated with different center frequencies and spectral widths with task performance and confirmed that PSD slopes were positively correlated with task accuracy for most center frequencies regardless of spectral width (Figure 3F). This suggests that when examining the role of spectral slope, most ranges centered between 20-50 Hz should give congruent results.

### Assessing information content through sample entropy

The relation between spectral slope and accuracy may be partially explained by the complexity of the underlying iEEG signal which may in turn suggest a higher capacity for processing information. However, while spectral slope is related to signal complexity (Keshner, 1982), it is not a direct measure. We therefore calculated sample entropy to quantify signal complexity of the iEEG trace, a measure that has previously been successfully used for discerning differences between EEG signals (Figure 4A; see Materials and Methods) (Catarino et al., 2011; Mizuno et al., 2010; Vaz et al., 2017). Sample entropy measures the predictability of a signal, is robust to low level noise and artifacts, and has been found to be more robust for shorter data lengths than other measures of entropy such as approximate entropy (Sokunbi, 2014; Yentes et al., 2013). Indeed, the complexity of two example iEEG signals is visible in the raw recording and reflected in the measured sample entropy (Figure 4B).

**Figure 4.**
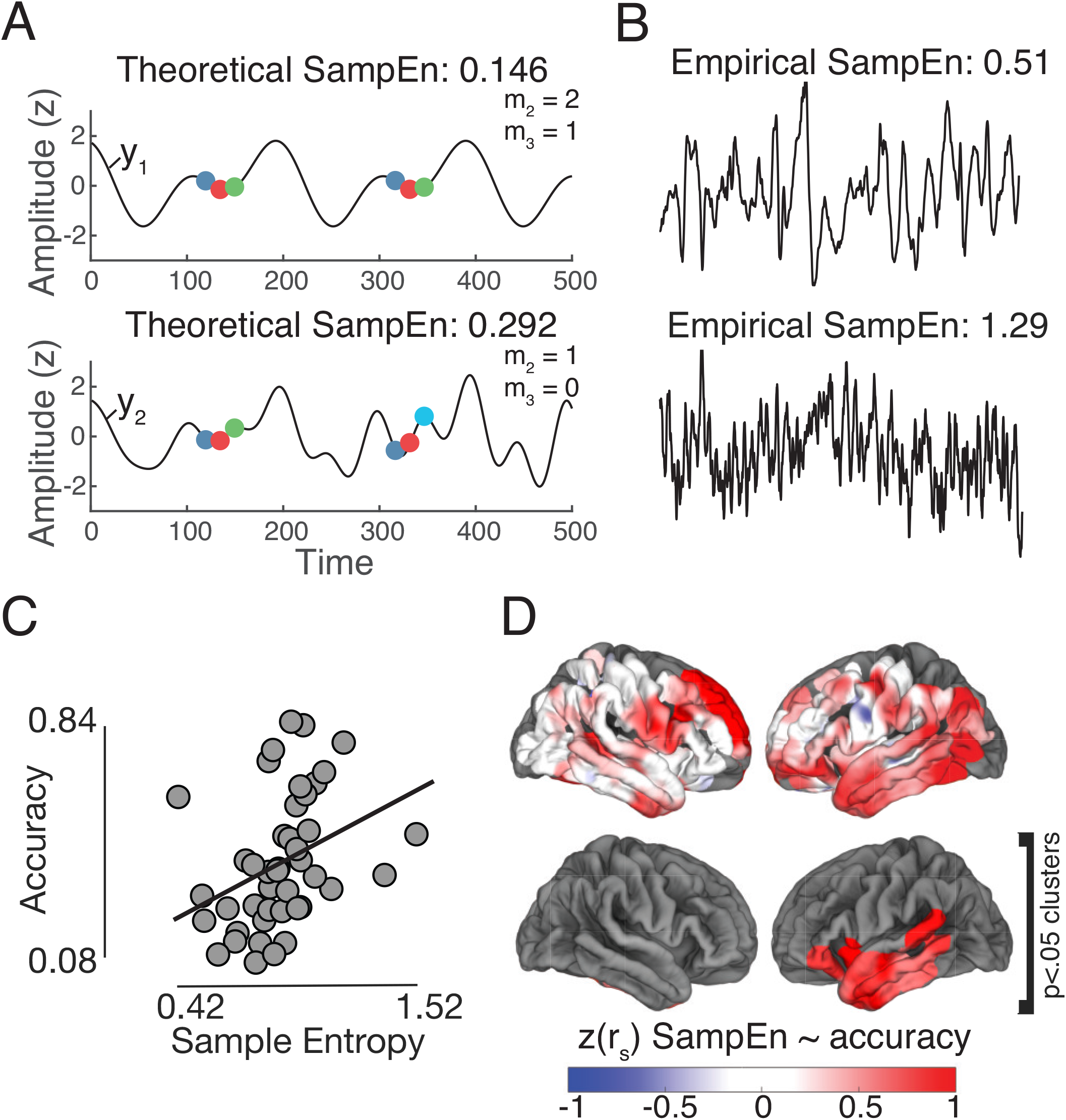
Sample Entropy and Performance. (*A*) Sample entropy schematic for theoretical signals. Color of dots superimposed on signals indicate discretized voltage bin. Signal *y*_2_ is more complex than *y*_1_ making subsequent points relatively more difficult to predict. (*B*) Example epochs from two participants with low and high entropy. The upper signal is from participant with an average sampEn of 0.51, this epoch has a measured sampEn of 0.67. The lower signal is from participant with an average sample Entropy of 1.29, this epoch has a measured sample Entropy of 1.51. (*C*) Sample entropy is positively correlated with performance across participants *r_s_* = 0.51, *p* = .0007. Line is standard least squares regression line. (*D*) Sample entropy correlation across spatial ROIs. Lower panel shows regions significant (*p* < .05) compared to a permuted distribution through a clustering procedure.

Based on the observed changes in raw power and spectral slope, and the theoretical suggestion that increased information content involves signal desynchronization (Hanslmayr et al., 2012), we hypothesized that participants with greater complexity in their iEEG signal, and therefore higher sample entropy, would exhibit better task performance. We calculated the average sample entropy during the encoding period across all trials and all electrodes and found that participants with greater sample entropy performed significantly better on the task (Figure 4C, *r_s_* = 0.51, *p* = .00065, *N* = 43), suggesting that the observed relation between task performance and PSD slope is related to the complexity of the underlying neural signal. Similar to slope, sample entropy was not correlated with IQ, (*r_s_* = 0.26, *p* = .14) and was correlated with accuracy after regressing out the effects of IQ (*r_s_* = 0.52, *p* = .0015, *N* = 35). The relation between sample entropy and accuracy was distributed across the cortex but was particularly localized to the left temporal lobe (Figure 4D).

As with power and spectral slope, we found that this relation was preserved when looking at correct trials (*r_s_* = 0.48, *p* = .0012), incorrect trials (*r_s_* = 0.52, *p* = .00049), the baseline period (*r_s_* = 0.53, *p* = .00028), or an extra-task epoch (*r_s_* = 0.34, *p* = .025). Moreover, participants with flatter PSD slopes and less theta power had greater sample entropy (*r_s_* = 0.63, *p* = 1.2 × 10 ^5^ and *r_s_* = –0.41, *p* = .0067, respectively) demonstrating that spectral slope and low frequency power are strong indicators of signal complexity.

### Signal complexity across time scales

Our data demonstrate that related measures of signal complexity — low frequency power, spectral slope, and sample entropy — show strong relations with overall performance during an associative memory task across participants. However, most studies of human memory have focused on subsequent memory effects in which differences between correct and incorrect trials are assessed within individuals. We were therefore interested in whether the observed changes in neural signal complexity across participants would also be observed across different time scales within participants. We specifically investigated changes in sample entropy during individual sessions and trials to index changes in brain state complexity at the time scales of hours and seconds, respectively.

We first examined the relation between sample entropy and performance during individual sessions for each electrode in each participant who completed at least three sessions. In individual participants, we found that sample entropy correlated with performance on a session by session basis (Figure 5A). Across all participants, we found that this relation was consistent, although the distribution of correlation coefficients was not significantly different from zero (two-tailed *t*-test of Fisher transformed correlation coefficients, *t*(21) = 2.00, *p* = .058; Figure 5B). However, while this relationship was not significant on a global level, we did find that individual ROIs throughout the left temporal and parietal lobes were significant (Figure 5C). In addition, we found that theta power and slope also showed strong relationships with session accuracy that were similar to those found across participants (*t*(21) = –2.37, *p* = 0.028 and *t*(21) = 1.83, *p* = 0.081, respectively).

**Figure 5.**
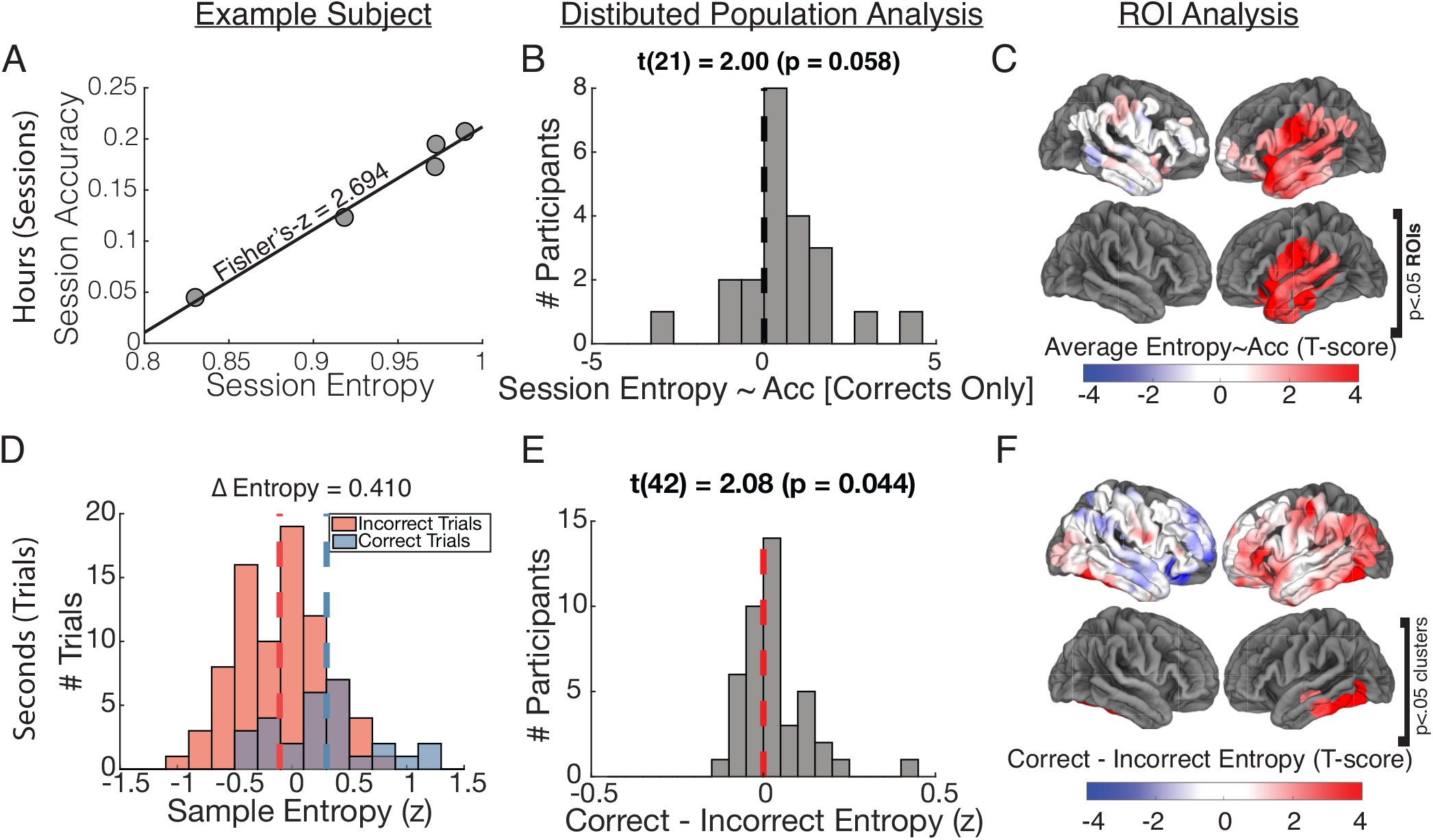
Sample Entropy across Time Scales. (*A*) Example subject with positive session level correlation of sample entropy to accuracy (*B*) Distribution of fisher transformed *ρ* values across subjects is trends towards positive correlations (*t*(21) = 2.00, *p* = .058). (*C*) Population average session correlation by ROI (*t*-score). Lower panel shows independently significant ROIs (*p* < .05, uncorrected) (*D*) Example subject level distribution of sample entropy values for correct vs. incorrect trials (*E*) Distribution of correct - incorrect sample entropy across subjects is significantly greater than 0 (*t*(42) = 2.08, *p* = .044) (*F*) Population average change in sample entropy by item (*t*-score) across ROIs. Lower panel shows regions significant (*p* < .05) compared to a permuted distribution through a clustering procedure.

Next, as is routine in most memory studies, we examined differences between correct and incorrect trials to understand the relation between sample entropy and memory encoding at the time scale of seconds. We first *z*-scored sample entropy within each session to eliminate any session level variance. Hence, both the session level and item level effects are calculated such that they are completely independent from one another and the previously explored participant level effects. We found that participants exhibited significantly higher sample entropy for correct compared to incorrect trials (Figure 5D,E, *t*(42) = 2.08, *p* = .0044). The item level changes in sample entropy localized to the left inferior temporal lobe (Figure 5E). The differences between correct and incorrect theta power and slope were also both significantly biased in the same direction as their across subject and across session effects (*t*(42) = –2.57, *p* = 0.014 and *t*(42) = 2.06, *p* = 0.046, respectively). To complement our analysis across participants, we also examined whether individual SMEs were correlated with raw power measurements. While slope and sample entropy showed no relationship (*p* > .90), raw theta power was significantly and negatively correlated with the difference between correct and incorrect theta power. Participants with greater raw theta power had more negative SMEs, while those with less raw theta power had more positive SMEs (*r_s_* = –0.33, *p* = 0.031). This suggests that, while by far the overriding trend in our cohort is that less theta power is better for encoding across all time scales, for a subset of participants with overall low theta power, this is not the case.

In evaluating across subject SMEs, it is unclear over what time scale the changes in sample entropy are occurring. These changes may be related to word pair adaptation, or they may reflect a more slowly fluctuating dynamic. To explore this, we made three additional comparisons. We compared the sample entropy during the baseline period between correct and incorrect trials, we compared the sample entropy during correct trials between the encoding and baseline periods, and we compared the sample entropy during incorrect trials between the encoding and baseline periods. Interestingly, while we found minimal difference in sample entropy during the baseline periods between correct and incorrect trials (*t*(42) = –1.71, *p* = .095), we did find that sample entropy significantly increased from baseline during correct trials, and significantly decreased from baseline on incorrect trials (*t*(42) = 2.41, *p* = .020 and *t*(42) = –2.36, *p* = .023). These data suggest that relatively fast changes in the sample entropy, and therefore complexity, of the signal contribute to subsequent remembering and subsequent forgetting along with changes over much longer longer time scales.

### Are theta power, spectral slope, and sample entropy redundant?

Sample entropy was shown earlier to be positively correlated with spectral slope and negatively correlated with theta band power. Spectral slope and theta band power are inversely correlated with each other as well *r_s_* = –0.35, *p* = .02. It is unclear, given the high level of collinearity between these variables, whether they are describing unique underlying processes or are really redundant factors. To determine the proportions of variance in memory performance across participants that are uniquely attributed to these metrics as well as those that are common between all possible combinations of these metrics, we performed a commonality analysis (see Materials and Methods). The commonality analysis (Table 1) showed that spectral power in the theta band uniquely accounted for 22.28% of the total variance explained by the predictors, with slope and entropy uniquely explaining 6.98% and 8.64%, and jointly explaining 20.04% of the total variance explained. The total variance accounted for by power, PSD slope, and sample entropy through both unique and shared contributions are 24.58%, 22.58%, and 25.52% respectively with 10.75% of the variance being common to all three. This analysis illustrates that, while all three metrics capture properties of neural activity that are relevant for memory performance, theta power may capture somewhat distinct features than those that are captured together by spectral slope and sample entropy.

**Table 1.** Commonality analysis output describing unique and common contributions of the three predictor variables (theta-band power, spectral slope, and sample entropy) to the regression effect explaining memory performance across participants.

|  | **power** | **spectral**<br>**slope** | **sample entropy** | **% Total** |
| --- | --- | --- | --- | --- |
| Unique to power | 0.0851 |  |  | 22.28 |
| Unique to spectral slope |  | 0.0267 |  | 6.98 |
| Unique to sample entropy |  |  | 0.0330 | 8.64 |
| Common to power and spectral slope | 0.0151 | 0.0151 |  | 3.96 |
| Common to power and sample entropy | 0.0381 |  | 0.0381 | 9.97 |
| Common to spectral slope and sample<br>entropy |  | 0.0766 | 0.0766 | 20.04 |
| Common to power, spectral slope, and<br>sample entropy | 0.1075 | 0.1075 | 0.1075 | 28.12 |

Total $R^2 = 0.3821$ . Unique + Common = 100% of $R^2$ .

To see how these metrics relate across time, we looked at their correlations across trials for individual participants. Across trials, *z*-scored by session, theta power and sample entropy showed a strong negative correlation (median *ρ* = –0.77). This aligns with our interpretation that, from an information coding perspective, predictable oscillations contain less information than less predictable stochastic dynamics. As expected, entropy and slope were positively correlated on a trial level (median *ρ* = 0.59) and slope and theta power were inversely correlated (median *ρ* = –0.60). Notably, these relationships were all highly significant on a population level (|*t*(42)| > 17, *p* < 10^−20^).

## Discussion

Our analyses demonstrate that low frequency power and complexity of cortical activity track variability in memory performance at the level of individuals, days, and events. Specifically, improved memory performance at any given scale is related to decreased low frequency power and increased signal complexity as measured by the PSD slope and sample entropy. Notably, while these metrics show collinearity on both an individual and population level, commonality analysis revealed substantial unique contributions of each metric in explaining memory performance. These findings suggest that the complexity of brain activity may reflect an individual’s ability to occupy variable cognitive states and the extent to which information can be coded in their brain signals, which is evident in associative memory performance.

The suggestion that cognitive flexibility may improve task performance appears intuitive. Indeed, the ability to explore the brain’s dynamic repertoire during rest is thought to be a marker of healthy brain function and may underlie introspection and rehearsal (Ghosh et al., 2008). Therefore, it seems likely that a high performing brain is one that engages with the world by assuming a variety of functional configurations. Whether such variability and flexibility may be relevant for associative memory performance has, until now, not been directly established. We establish this link here by demonstrating that memory performance is significantly correlated with signal complexity both across and within individuals. Cognitive flexibility lends neural systems the ability to explore their state space (Deco and Jirsa, 2012) which may lead to separable memory representations that are less susceptible to interference. Consistent with the idea that increased complexity may lead to increased separability of events, pre-stimulus weighted permutation entropy of scalp EEG can bias participants’ perception of identical auditory stimuli by changing the fidelity with which the stimulus was encoded (Waschke et al., 2017). Furthermore, multiscale entropy (MSE) of brain signals (scalp EEG) correlate with participants’ ratings of famous face familiarity and increase with learning over multiple exposures to previously unfamiliar faces (Heisz et al., 2012). Hence, the observed correlations between entropy and associative memory performance here suggests that neural signal complexity reflects the capacity to successfully encode associative memories by flexibly engaging with the presented material.

The paired associates memory task used here requires participants to form associations between unrelated words that constitute individual episodes or experiences that are subsequently recalled. Encoding these associations draws upon the meanings of the words in order to form a conceptual and semantic link between them (Jang et al., 2017; Kahana et al., 2008; Madan et al., 2010). Therefore, forming these associations should engage cortical regions such as the anterior temporal lobe that are involved in semantic processing (Binder et al., 2009; Ralph et al., 2017). In our data, we observe strong correlations between memory performance and low frequency power and entropy in these same left temporal lobe regions. The relationship between cognitive flexibility, as assessed by these metrics in the temporal lobe, and verbal associative memory performance across individuals may therefore emerge because of the involvement of the temporal lobe in helping encode verbal associations.

Our approach here differs from earlier studies of human memory encoding and retrieval by specifically asking whether there are systematic differences in neural activity across participants that may predict individual memory performance. Most previous studies of human episodic memory have focused on relative changes in neural oscillatory activity between correctly and incorrectly encoded events (Burke et al., 2014; Greenberg et al., 2015; Long et al., 2014; Sederberg et al., 2003, 2007). While these studies have significantly advanced our understanding of the neural correlates of human memory, an unresolved question has been why different studies have demonstrated conflicting results, particularly with respect to low frequency oscillatory power (Hanslmayr and Staudigl, 2014). In our data, we tracked memory performance using low frequency power that was not normalized relative to any baseline and found that it was inversely correlated with overall memory performance. Moreover, within each individual, fluctuations in neural activity were predictive of how well they performed at any given moment. Interestingly, we found that these fluctuations were dependent on overall power measurements for each participant. Our data therefore may provide some insight into the conflicting data observed in previous studies. These conflicts have been previously attributed to differences in task design and electrode coverage. However, because of the variability in baseline power between individuals, these conflicts may also be affected by where each cohort of participants sits in this range of baseline power and how that may impact the changes in power observed over shorter time scales.

As examining the structure of the full PSD across all frequencies can often yield a more complete picture of neural activity (Podvalny et al., 2015), our analyses of PSD slope changes complement the observed changes in low frequency power. The slope of the PSD has been hypothesized to reflect the balance between excitation and inhibition, and computational modeling of neural activity has demonstrated that reducing E:I ratio results in a steeper PSD slope (Gao et al., 2017). Both *in vitro* and *in vivo* cortical networks show maximal dynamic range under balanced E:I conditions (Shew et al., 2009, 2011). An increased dynamical range of neuronal responses may improve adaptability and efficiency of neural systems in service of memory. Another possibility is that a shallower PSD slope may emerge due to the infusion of noise into the neural signal via asynchronous firing activity (Podvalny et al., 2015; Pozzorini et al., 2013; Usher et al., 1995; Voytek et al., 2015; Voytek and Knight, 2015). Whether such noise is beneficial is unclear, as the effect of noise on information coding depends on whether or not noise is correlated between neurons (Averbeck et al., 2006). Nevertheless, our finding that flatter PSD slopes and increased sample entropy relate to better memory performance suggests that in our data, more complex brain signals reflect more informationally rich signals as posited by others (Hanslmayr et al., 2012, 2016; Mitchell et al., 2009; Schneidman et al., 2011).

Ultimately, the slope of the PSD and the low frequency power contributing to that slope should be related to the underlying complexity of the neural signal, which can be directly assessed using measures of entropy as we do here. Although greater signal complexity does not always reflect greater information content, entropy of the EEG signal may increase with healthy aging (McIntosh et al., 2008; Waschke et al., 2017), and higher entropy is also associated with greater task efficiency and network efficiency (Misić et al., 2010, 2011). Entropy of resting state brain signals can distinguish children at high risk for autism spectrum disorder from normal developing children (Bosl et al., 2011), and healthy from epileptogenic neural tissues (Protzner et al., 2010). Here, we directly show that the complexity of the neural signal captured using iEEG tracks associative memory performance across individuals, providing further support to the proposition that brain signal variability is functionally relevant. Moreover, we show that within individuals, variability in neural signal complexity across time scales also tracks memory performance for the individual participant. The variability that we experience in our daily lives with memory performance is likely therefore influenced by these changing levels of neural signal complexity.

Of note, participants in our study were also neurosurgical patients with drug resistant epilepsy. In the majority of cases, their seizure activity localized to the temporal lobes, raising the possibility that the observed effects in this brain region may also be related to the underlying pathology of the disorder itself. Greater disruptions of normal temporal lobe function could result in less signal complexity in this brain region, which could then lead to worse memory performance on this paired associates task. We took several precautions to mitigate the effects of epileptic activity on our study including removing electrodes identified as ictal or interictal, and removing electrodes and individual trials that showed higher variance or kurtosis relative to the rest of the population. In addition, we visually checked for both high and low amplitude IEDs in electrodes and trials chosen from a subset of participants. While we found that IEDs were approximately equally present in both high and low power trials and electrodes, it is also clear that our data were not devoid of these artifacts. Hence, it is possible that pathological activity may contribute to some of the observed relationships between complexity metrics and memory performance. In this scenario, however, the interpretation of our data does not change, since decreased neural signal complexity, regardless of whether it can be attributed to normal or pathologic variability, would still be related to decreased memory ability

Previous studies have indeed shown that interictal epileptiform discharges (IEDs) during encoding and retrieval can impair memory performance (Horak et al., 2017), and IED rates decrease from baseline during correct, but not incorrect, encoding trials (Matsumoto et al., 2013). Critically, however, increases in IEDs during rest or distractor periods in these studies do not appear to reduce memory performance, and the overall IED rates do not relate to recall performance across participants (Horak et al., 2017). This is in contrast to the effects of overall power on memory performance that we report here, which are observed during both rest and task periods. Moreover, controlling for the same level of overall pathology within individuals, we find the same metrics were related to memory performance across multiple timescales. It is difficult to explain how pathologic activity alone would consistently predict memory performance at every different timescale, or even why most effects in our data also extend to generally non-pathologic frontal lobe clusters in our data set.

In addition to the brief disruptions in temporal lobe function caused by interictal epileptiform discharges, transient neurologic dysfunction can be observed after a seizure lasting minutes to hours. In the case of left temporal lobe epilepsy, post-ictal impairment can be seen in verbal and visual recognition memory. However, post-ictal effects are unlikely to play a significant role in our findings, since we avoided administering cognitive testing for several hours following a seizure episode. In addition, if patients were still symptomatic following a seizure, testing was usually deferred by the study team or by the participant until they had regained their baseline function. The longer term effects of such pathologic activity may, however, contribute to changes in IQ, which could in turn mediate the across-participant relationship between signal complexity and memory. However, we found that only theta power is significantly correlated with IQ. Furthermore, theta power, PSD slope, and sample entropy are all significantly correlated with performance even after the effects of IQ are regressed out, suggesting that signal complexity is indeed specifically relevant for memory formation. Therefore, although the participants’ underlying disorder may affect normal neural information processing and overall cognitive ability, our data suggest that the individual differences in neural signal complexity that relate to differences in memory performance are unlikely to be driven by pathology alone.

It is also possible that the changes in neural complexity that we interpret to denote cognitive flexibility in fact simply capture changing levels of attention. For example, patients may feel drowsier in some experimental sessions than others and these differences in levels of engagement may be captured by our complexity metrics. However, we note that changes in sample entropy from baseline to encoding states also occur over shorter timescales within an individual experimental session. These fine-grained temporal changes consistently capture differences between successful and unsuccessful associative memory encoding trials even though the baseline entropy levels are not different between the two conditions. Moreover, at the other extreme of time scales, participant-level complexity metrics correlate with memory performance as well as IQ. Therefore, it is unlikely that drowsiness explains all of the observed relationships found here between neural complexity and memory performance at multiple scales. Attention may indeed play a direct role in determining the extent to which neural state space is explored during a task. However, the possibility that changing levels of attention may explain our results is still consistent with the interpretation that theta power, spectral slope, and sample entropy ultimately reflect cognitive flexibility and a capacity to encode information.

Aside from the immediate effects of interictal and ictal epileptiform activity, it is also possible that some of these relationships are impacted by the influence of anti-epileptic drugs (AEDs). All of the participants were chronically taking AEDs, which were weaned at a variable rate following surgery at the discretion of the treating clinicians. Because participants were on varying AEDs at varying doses with varying pharmacokinetics, we did not explicitly control for AEDs as a factor in our study. In general, since testing began several days post-operatively, AEDs were at a significantly lower level than at baseline for a given participant. AEDs are known to reduce attention and vigilance, but other studies have suggested that the cognitive impacts of AEDs may be overrated when compared to the pathological and psychosocial effects of epilepsy itself (Meador, 2002; Park and Kwon, 2008). Studies exploring electrophysiological changes related to AED use have found highly heterogenous results across participants and medications, with some evidence for short term reductions in gamma power (Arzy et al., 2010) and long term slowing of EEG rhythms (Salinsky et al., 1994), measures that have minimal overlap with our metrics. Hence, while AEDs may certainly be a relevant factor, they are unlikely to be primary driving force of the reported effects that persist across timescales and individuals.

Together, our data therefore provide insight into why memory performance may be variable both between and within individuals. Our data suggest that how well one can encode and retrieve memories is related to the flexibility in their cognitive processing. Such flexibility is captured directly by measuring the sample entropy of the neural signal, and corroborated by our measures of low frequency power and the PSD slope. People with better memory have neural signals that exhibit greater complexity, and therefore are capable of exhibiting more flexible behavior that is beneficial for memory formation.

## Acknowledgements

We thank John Wittig Jr. and Julio Chapeton for helpful comments on the manuscript. We are indebted to all patients who have selflessly volunteered their time to participate in this study. This work was supported by the Intramural Research Program of the National Institute for Neurological Disorders and Stroke. This work was also partially supported by the DARPA Restoring Active Memory (RAM) program (Cooperative Agreement N66001-14-2-4032). The views, opinions, and/or findings contained in this material are those of the authors and should not be interpreted as representing the official views or policies of the Department of Defense or the U.S. Government.

The authors declare no competing financial interests.

